# Specificity in auxin responses is not explained by the promoter preferences of activator ARFs

**DOI:** 10.1101/843391

**Authors:** Amy Lanctot, Mallorie Taylor-Teeples, Erika A. Oki, Jennifer L. Nemhauser

## Abstract

Auxin is essential for almost every developmental process within plants. How a single small molecule can lead to a plethora of downstream responses has puzzled researchers for decades. It has been hypothesized that one source for such diversity is distinct promoter-binding and activation preferences for different members of the AUXIN RESPONSE FACTOR (ARF) family of transcription factors. We systematically tested this hypothesis by engineering varied promoter sequences in a heterologous yeast system and quantifying transcriptional activation by ARFs from two species, *Arabidopsis thaliana* and *Zea mays*. By harnessing the user-defined and scalable nature of our synthetic system, we elucidated promoter design rules for optimal ARF function, discovered novel ARF-responsive promoters, and characterized the impact of ARF dimerization on their activation potential. We found no evidence for specificity in ARF-promoter interactions, suggesting that the diverse auxin responses observed in plants may be driven by factors outside the core auxin response machinery.

## Introduction

Promoter architecture is a key determinant of specificity in the activation of downstream genetic networks. Animal steroid hormone receptors are perhaps the best-understood model for how a common ancestral transcription factor can diverge to produce multiple proteins with high selectivity for distinct promoter sequences (McKeown et al., 2014). In plants, hormone response is essential to plant growth and development and also involves large gene families, particularly in the auxin response. Whether a similar evolutionary trajectory is at work in the auxin response has been a long-standing question. When auxin enters the nucleus, AUXIN/INDOLE-3-ACETIC ACID (Aux/IAA) co-repressor proteins are degraded, relieving repression on AUXIN RESPONSE FACTOR (ARF) transcription factors and allowing them to induce the transcription of downstream genes (Chapman and Estelle, 2009). It has been hypothesized that different ARFs bind to and activate on distinct promoters and that this is how an auxin signal can lead to a diversity of transcriptional responses (Boer et al., 2014; O’Malley et al., 2016).

ARFs are comprised of large gene families in most angiosperms (Remington et al., 2004), and the subset of ARFs that activate transcription likely do so through multiple mechanisms. The AUXIN RESPONSE FACTOR (ARF) family of transcription factors has 23 members in *Arabidopsis*, five of which are classified as activators (Guilfoyle and Hagen, 2007). *Zea mays* has 33 expressed ARFs, thirteen of which cluster with the activator clade in *Arabidopsis* (Galli et al., 2015). AtARF5 has been shown to recruit chromosome-remodeling ATPases to change nucleosome occupancy on actively transcribed promoters (Wu et al., 2015), and AtARF7 and AtARF19 can interact with Mediator subunits (Ito et al., 2016). ARFs bind DNA as dimers and loss of dimerization leads to decreased DNA binding (Boer et al., 2014) and activity (Pierre-Jerome et al., 2016).

While the activator ARFs are co-expressed within many cells (Rademacher et al., 2011), they have distinct developmental roles (Krogan et al., 2016; Wilmoth et al., 2005). For example, AtARF5 regulates embryonic and primary root development (Hardtke and Berleth, 1998; Schlereth et al., 2010) while AtARF7 and AtARF19 regulate lateral root development (Okushima et al., 2005; Okushima et al., 2007). These distinct roles may be mediated by differing promoter preferences among the ARFs, allowing them to activate different target genes. ARFs bind to the <u>aux</u>in-responsive cis-<u>e</u>lement, or AuxRE. This sequence was first described in *Pisum sativum* as the six-mer TGTCTC/GAGACA (Ballas et al., 1993); however, further work revealed that there is some flexibility in the fifth and sixth base pairs. Though all activator ARFs can bind to the canonical AuxRE sequence *in vitro*, promoter context may allow for specificity in ARF-promoter interactions *in vivo*. For instance, auxin response in several *Glycine max* promoters requires an AuxRE but additionally require an upstream constitutive activation sequence, suggesting that surrounding sequences can influence both auxin-inducibility and strength of transcriptional response (Ulmasov et al., 1995).

Several recent studies that focus on cross-clade comparisons, particularly between the Class A (“activator”) and Class B (“repressor”) ARFs, support a model of ARF-specific binding preferences. High-resolution crystal structures of ARF DNA-binding domains and *in vitro* binding assays suggest that AtARF5 (Class A) and AtARF1 (Class B) homodimers exhibit different stringency in the numbers of nucleotides between pairs of binding sites (Boer et al., 2014) on which they can activate. Similar results are reported in DAP-seq data in maize and *Arabidopsis*, which reveal distinct spacing and orientation preferences for Class A versus Class B ARFs (O’Malley et al. 2016, Galli et al., 2018). While the DAP-seq studies have led to a wealth of information on ARF binding, their analytical power is limited to the variation found within native genomes. In addition, DAP-seq clusters a large number of DNA fragments according to investigator hypotheses about functional features, leaving open the possibility that differences in promoter structure are missed. Another complication in interpreting these data is that transcription factor binding to DNA and activation at a given locus are often decoupled (Para et al., 2014).

To complement these ARF binding studies, we tested the activation profile of Class A ARFs from *Arabidopsis* and maize on synthetic, user-defined promoter sequences using a heterologous yeast activation system (Pierre-Jerome et al., 2014). This approach allowed us to test the hypothesis that the observed differences in transcriptional profiles induced by different ARFs might reflect differences in ARF activity on distinct promoters. We conducted our assays in a pairwise fashion, looking at each ARF-promoter interaction individually, on standardized promoter variants to directly test how of promoter architecture affects activity. The synthetic system also allows us to survey a sequence space unreachable by *in planta* studies that are limited to native promoters. We queried the activity of two subclades of Class A ARFs, the AtARF5 clade (ZmARF4 and ZmARF29) and the AtARF19 clade (ZmARF27). We found that Class A clade ARFs across species largely shared promoter preferences, and additionally found that AtARF19 was the only ARF tested to be able to activate transcription on promoters with a single AuxRE. Promoter preferences were shared across subclades of ARFs as well as conserved between *Arabidopsis* and maize.

## Results

### Class A ARFs prefer similar promoter architectures in terms of cis-element number and orientation

A long-standing question in the field of auxin biology is how different members of the ARF gene family regulate different genes. Several studies have shown that ARFs bind to and activate on different promoter sequences to varying degrees (Boer et al., 2014; Pierre-Jerome et al., 2016), giving rise to the hypothesis that ARF-promoter interactions may lead to specificity in downstream response. We used flow cytometry on engineered yeast to test how Class A ARFs from two clades, the AtARF5 and AtARF19 clades, activate on synthetic promoter variants (Figure 1A). All sequence variants were embedded within the same genomic context: the first 300-base pairs of the *Arabidopsis thaliana* IAA19 promoter with all five putative auxin responsive elements (AuxREs) mutated (mpIAA19). None of the ARFs tested can activate transcription to any appreciable extent on the mpIAA19 promoter (Supplemental Figure S1). Variants were specifically embedded at the A1 position, an AuxRE 166 base pairs from the transcriptional start site (TSS) (Pierre-Jerome et al., 2016). This region relative to the TSS has been shown to be enriched for AuxREs within the *Arabidopsis* genome (Lieberman-Lazarovich et al., 2019).

**Figure 1.**
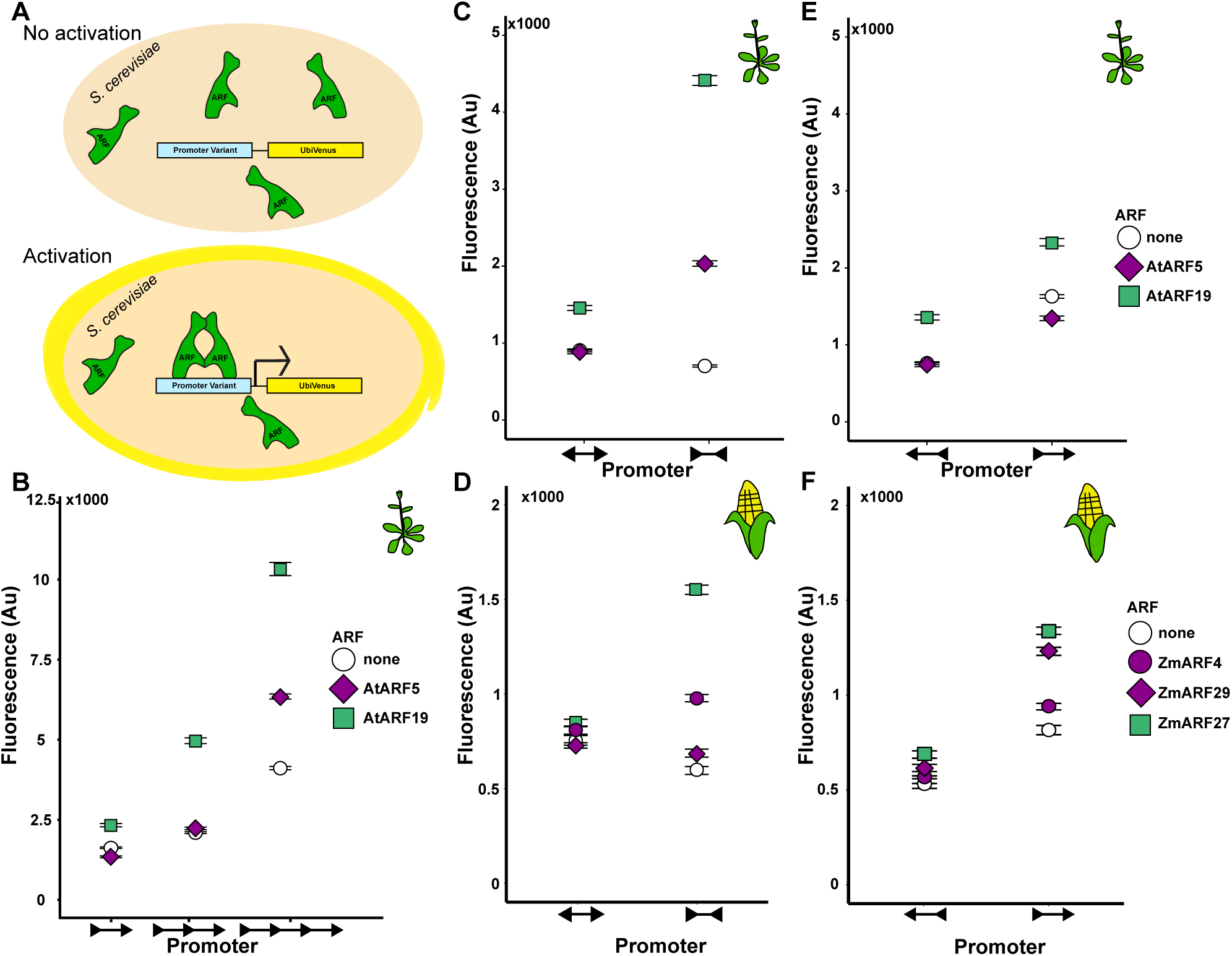
Arabidopsis and maize ARFs share promoter preferences. A) Schematic of yeast engineered to constitutively express ARF proteins and promoter variants. All promoter variants were inserted into the A1 site of a pIAA19 promoter with mutated AuxREs. The transcription start site (TSS) is to the right and arrowheads indicate the orientation of the AuxRE, starting with 5’-TGTC-3’. Fluorescence was measured by flow cytometry with the results depicted as median values and 95% confidence intervals. B) AtARF19 and AtARF5 show strong activation on promoters with four AuxREs (five base pair spacer). C) AtARF19 and AtARF5 show stronger activity on promoters with two AuxREs facing towards each other rather than away from each other (seven base pair spacer). D) ZmARF4, ZmARF27, and ZmARF29 show stronger activity on promoters with two AuxREs facing towards each other rather than away from each other (seven base pair spacer). E) AtARF19 and AtARF5 show stronger activity on promoters where the two AuxREs face towards rather than away from the TSS (five base pair spacer). F) ZmARF4, ZmARF27, and ZmARF29 show stronger activity on promoters where the two AuxREs face towards rather than away from the TSS (five base pair spacer).

We first tested how the copy number of AuxREs within a promoter affects activation by AtARF5 and AtARF19. We generated three copy number variants, with two to four copies of the canonical forward-facing AuxRE TGTCTC. A five base pair spacer CCTTT separated these AuxREs, which is the spacer sequence in the commonly used auxin-responsive DR5 reporter (Ulmasov et al., 1997b). We found that the activation strength of both AtARF5 and AtARF19 was directly proportional to AuxRE copy number, with the highest activation by both ARFs on the promoter with four AuxREs (Figure 1B). It is worth noting that AtARF5 activation was significantly lower than that of AtARF19, making it difficult to assess whether it was able to activate at all on promoters with less than four AuxREs, and that background activity also increases with increased AuxRE copy number.

We next tested how the orientation of AuxREs relative to each other and to the TSS affects activation. For this we generated two sets of two promoter variants (four total) all containing two AuxREs. In the first set, we tested whether ARFs activated more strongly on two AuxREs facing towards each other, separated by seven base pairs, or two AuxREs facing away from each other, separated by the same seven base pair sequence. We used the canonical AuxRE sequence TGTCTC and the spacer sequence from the ER7 auxin reporter, CCAAAGG. We found that all the tested ARFs activated more strongly on two AuxREs facing towards each other rather than away from each other (Figures 1C and 1D), and neither AtARF5 nor the tested ZmARFs showed appreciable activation when the two AuxREs were facing away from each other compared to the background yeast activation.

We also examined AtARF5 and AtARF19 activation on two promoters with two AuxREs facing either towards the TSS or away from the TSS. In these promoter variants the AuxREs were spaced by five nucleotides, and the spacer sequence was the one used previously in DR5 reporters. We found that AtARF19 activated slightly more strongly when AuxREs face towards the TSS as opposed to away from the TSS (Figure 1E). AtARF5 did not activate on two AuxREs facing either towards or away from the TSS when compared to background yeast activation, indicating that AtARF5 is a weaker activator than AtARF19. None of the ZmARFs strongly activate on two AuxREs facing away from the TSS, while ZmARF27 and ZmARF29 activate to some degree on two AuxREs facing towards the TSS (Figure 1F). Interestingly, this is the only orientation on which ZmARF29 appreciably activated. Of note, background activation increases on two AuxREs facing towards the TSS, but comparison to a control strain of yeast expressing no ARFs allows the determination of ARF-dependent activation. All of the ARFs we tested activated most strongly on two AuxREs facing towards each other, and activated weakly or not at all on two AuxREs in any other orientation. This is the orientation for the solved structures of the AtARF5 and the Class B AtARF1 DNA-binding domains (Boer et al., 2014).

### AtARF5 more strongly activates on the AuxRE TGTCGG than the canonical cis-element TGTCTC

While the canonical AuxRE is widely considered to be the TGTCTC and its reverse complement GAGACA, the “core” element is TGTC/GACA and auxin responsiveness has been seen on a wide variety of cis-elements with varying base pairs in the fifth and sixth positions. AtARF1 and AtARF5 in fact bind most strongly to the AuxRE TGTCGG and its reverse on two AuxREs facing towards each other (Boer et al., 2014). Additionally, DR5 reporters using different AuxRE sequences showed variable activation in a transient expression assay (Lieberman-Lazarovich et al., 2019). We tested how AuxRE sequence impacts activation by AtARF5 and AtARF19 on two AuxREs facing towards each other by comparing activation on the AuxREs TGTCTC/GAGACA and on the AuxREs TGTCGG/CCGACA. We found that all tested ARFs activate more strongly on the TGTCGG/CCGACA AuxREs (Figures 2A and 2B). The difference in AtARF5 activation on the canonical AuxRE sequence and the novel sequence, nearly a nine-fold increase, was striking. In combination with previous protein binding microarray data (Boer et al., 2014), this may suggest AtARF5 has a strong preference for activation on TGTCGG/CCGACA, at least with this promoter orientation and spacer. Similarly, while the maize ARF5-like protein ZmARF4 does not activate well on TGTCTC/GAGACA, it does show transcriptional activity on the TGTCGG/CCGACA AuxREs at levels similar to ZmARF27. These results again do not show divergent promoter preferences among ARFs—while the relative degree of preference may differ between ARFs, they all activate more strongly on the same promoter variant.

**Figure 2.**
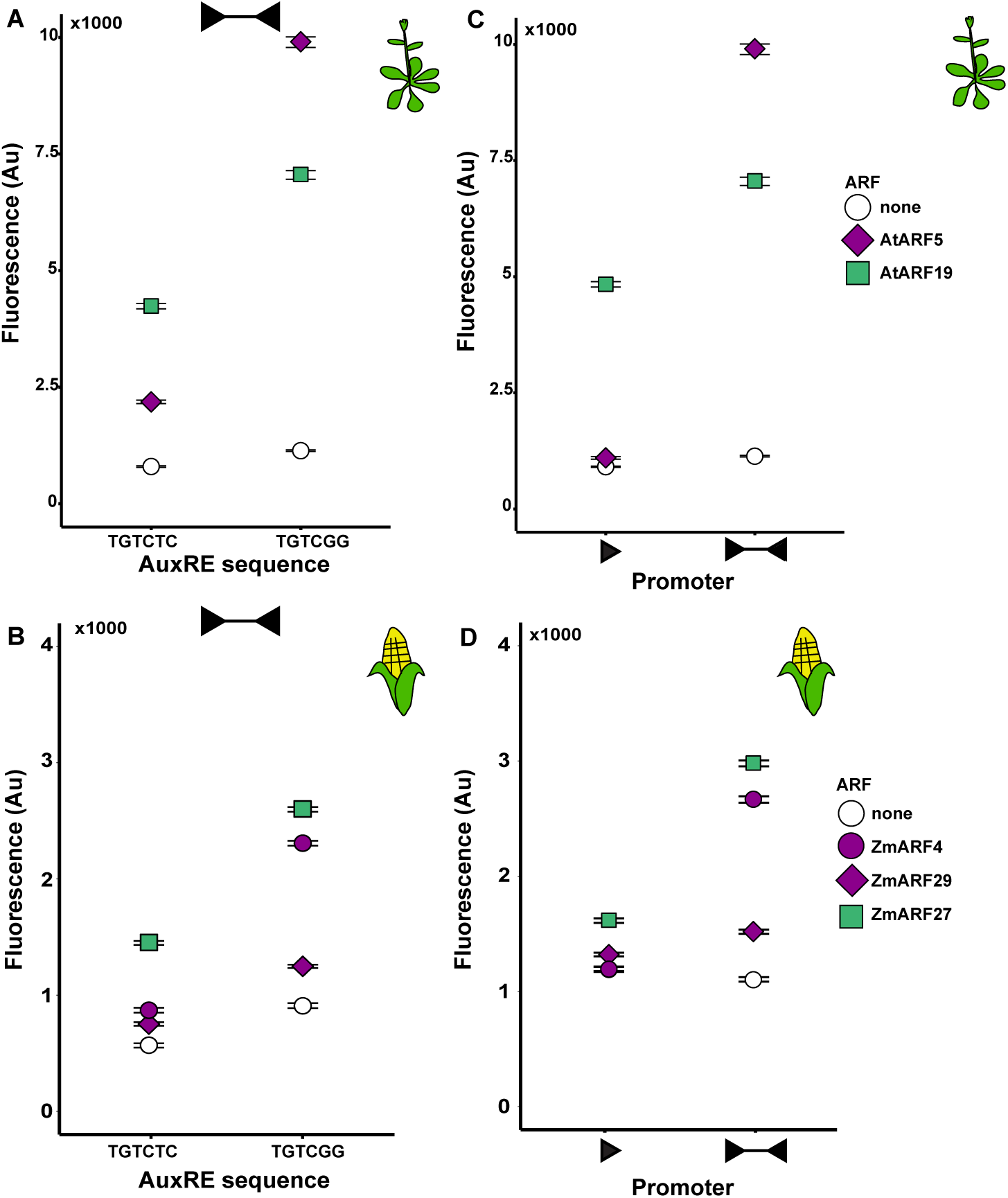
AtARF19 can activate on a single AuxRE of the sequence TGTCGG. A) AtARF5 and AtARF19 activate more strongly on two AuxREs facing each other of the cis-element sequence TGTCTC/GAGACA when compared to two AuxREs facing each other of the cis-element sequence TGTCGG/CCGACA. B) AtARF19, but not AtARF5, can induce transcription on a promoter with one AuxRE of the sequence 5’-TGTCGG-3’. C) ZmARF4, ZmARF27, and ZmARF29 activate more strongly on two AuxREs facing each other of the cis-element sequence TGTCTC/GAGACA when compared to two AuxREs facing each other of the cis-element sequence TGTCGG/CCGACA. D) None of the tested ZmARFs activate on a single AuxRE with the cis-element sequence 5’-TGTCGG-3’ (The no ARF control data point is directly underneath the ZmARF4 data point).

### AtARF19 can activate on a single AuxRE in yeast

Our results suggested that the AuxRE sequence TGTCGG and its reverse complement may be more optimal than the canonical AuxRE for ARF activation on the promoter. While common synthetic auxin responsive reporters have high copy numbers of AuxREs within a short sequence, in native auxin responsive promoters it is rare for two AuxREs to occur close together (Grigolon et al., 2018). To test whether ARFs can activate on a single AuxRE we placed the single AuxRE TGTCGG into the A1 site of the mutated pIAA19 promoter. Previous work from our lab showed that *Arabidopsis* ARFs cannot activate on a single AuxRE sequence that is natively in this position in the IAA19 promoter (TGTCGA) (Pierre-Jerome, 2016). To our surprise, we found that only AtARF19 was able to activate on this single AuxRE (TGTCGG) (Figures 2C and 2D). In fact AtARF19 activated almost as strongly on this promoter as it did when there were two TGTCGG AuxREs.

### Dimerization is required for ARF activity on single AuxRE promoters

ARFs have two dimerization domains, one at the N-terminus flanking the DNA-binding domain (termed the DD) (Boer et al., 2014) and one at the C-terminus (a Phox and Bem1 or PB1 domain) (Korasick et al., 2014, Nanao et al., 2014). Structural studies indicate that ARFs require dimerization at the DD to bind to DNA (Boer et al., 2014). In addition, mutations in either DD or PB1 of AtARF19 reduce ARF activity (Pierre-Jerome, 2016), though these studies only addressed ARF behavior on promoters with multiple AuxREs. We tested the activity of AtARF19 mutations that disrupt ARF dimerization in either the DD (G247I and A50N) or the PB1 domain (termed ARF19 KO—a triple mutation K962A; D1012A; D1016A) (Pierre-Jerome, et al. 2016) and compared these to the activity of a DNA-binding mutant AtARF19 H138A (Figure 3A, B). The dimerization mutations caused a loss of activation on the single AuxRE (TGTCGG) promoter to nearly the same extent as the DNA-binding mutation (Figure 3C), suggesting that dimerization is necessary for ARF activation on the promoter despite the presence of only a single optimal binding site. Interestingly, when we tested the activity of these dimerization mutants on the two TGTCGG AuxREs facing towards each other, they caused a loss of activation but not to the same extent as on the single AuxRE, suggesting that multiple AuxRE sites may compensate for a loss of dimerization of the ARFs themselves. As ARFs were crystallized as a dimer pair with each monomer bound to a separate AuxRE (Boer et al., 2014), how an ARF dimer contacts the DNA when there is a single AuxRE present is unknown. It is possible that only a single ARF-AuxRE interaction is required to bring the dimer to the DNA, and the other ARF forms transient interactions with multiple DNA sequences, which may serve as cryptic, low-affinity binding sites. Or the proximity of ARFs within a dimer pair may allow one to bind a single AuxRE promoter as soon as the other falls off, increasing the on rate of ARF binding to the promoter.

**Figure 3.**
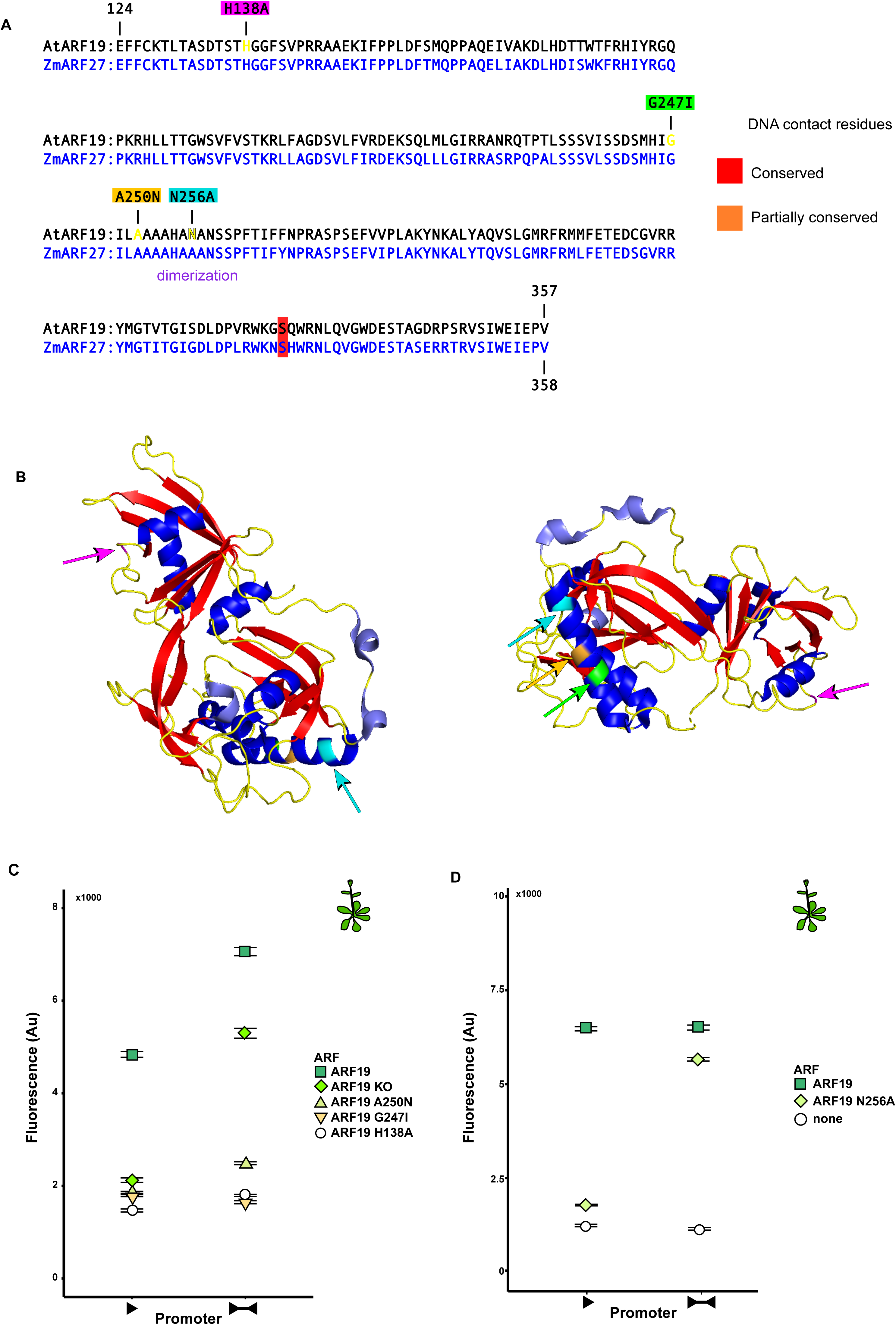
AtARF19 requires dimerization to activate even on a single AuxRE. A) Alignment of the DNA-binding and dimerization domains of AtARF19 and ZmARF27 with relevant mutations highlighted. B) Structure of ARF5 DNA-binding domain with mutated residues highlighted. C) AtARF19 must dimerize for full activity, even for a promoter with a single AuxRE. The KO mutation disrupts dimerization in the PB1 domain. The A250N and G247I mutations disrupt dimerization at the DD domain, adjacent to the DNA-binding domain. The H138A mutation disrupts the DNA-binding domain itself. D) An N256A mutation in AtARF19 causes a total loss of activity on a promoter with one AuxRE (5’-TGTCGG-3’), while leaving activity on two AuxREs largely intact.

### AtARF19 has a unique residue in the dimerization domain required for activity on a single AuxRE

Alignments among *Arabidopsis* and maize ARFs (Figure 3A) showed a difference in sequence within the DD of AtARF19 when compared to its maize homologues ZmARF27 and ZmARF35 (Figure 3B). We hypothesized that this single residue difference, so close to the monomer-to-monomer contact residues within the DD, could explain AtARF19’s unique ability to activate transcription on promoters with only a single AuxRE. To test this, we generated a mutated form of AtARF19 that replaced the asparagine residue with an alanine, the same amino acid found in ZmARF27 (N256A). This single residue change abolished AtARF19 activity on a single AuxRE, while leaving its activity on a two-AuxRE promoter essentially unchanged (Figure 3D). The polarity of the asparagine may help stabilize the dimeric form of AtARF19, leading to higher transcriptional activation overall and greatly increasing the number of potential promoters it can act on. While N256 is necessary for AtARF19’s ability to activate on promoter with a single AuxRE, it is not sufficient. AtARF7, which shares the same asparagine residue in its DD, cannot activate on a single AuxRE (Supplemental Figure S2). This difference, in combination with the critical role of the PB1 domain in ARF transcriptional activation (Figure 3C), implicates the still poorly understood inter-domain interactions in determining overall protein function.

## Discussion

It has been widely speculated that specificity within ARF-promoter interactions is responsible for the observed diversity in transcriptional and developmental responses triggered by auxin. Our results suggest that this model is unlikely to be true, at least among Class A ARFs. All the ARFs tested showed similar promoter preferences, and all required dimerization for full activity. We were able to elucidate a set of promoter design rules for maximizing response across the A clade, and found that these design rules were conserved across *Arabidopsis* and maize. Simply stated, these rules are as follows: (1) ARFs most strongly activate on promoters with at least four AuxREs arranged facing towards one another (Figure 1); (2) the non-canonical TGTCGG sequence can further boost expression, especially by ARFs in the AtARF5 clade (Figure 2). This second rule has relevance for the design and interpretation of auxin reporters. For example, DR5v2, which uses TGTCGG (Liao et al., 2015), may over-report responses driven by AtARF5 and its homologues relative to other Class A ARFs. Our study also highlights the complexity of inter-domain interactions within the ARFs, as dimerization at both N- and C-terminal dimerization domains was found to be critical for maximal transcriptional activation.

The differences between the architecture of auxin reporters and native auxin responsive promoters are striking. The rules derived from the systematic analysis presented here are generally consistent with the construction of auxin reporters, where there is a trend towards high copy numbers of canonical AuxREs in a short sequence space (Ulmasov et al., 1997a; Ulmasov et al., 1997b). Closely spaced AuxREs are found only rarely in the *Arabidopsis* genome (Grigolon et al., 2018), and frequently are neither the ideal sequence nor in the ideal orientation relative to the TSS. One possible explanation for the rarity of “ideal” auxin promoters is that it allows for integration of signals from multiple pathways, a hypothesis supported by the enrichment for transcription factor binding sites for other proteins in auxin-responsive promoters.

Our results showed that heterodimerization between ARFs is essential for ARF function, but importantly heterodimerization between ARFs and *other* transcription factors could support ARF activity on non-ideal native promoters and potentially act as a locus for specificity within auxin response. Bioinformatics analyses of auxin-induced genes show that many promoters of these genes are enriched for the binding sites of transcription factors such as bZIPs and bHLHs (Berendzen et al., 2012; Cherenkov et al., 2018; Mironova et al., 2014). Genetic studies show that heterodimerization between specific ARFs and members of other transcription factor families is required for the development of many plant organs, including lateral roots (MYBs; Shin et al., 2007) leaves (bHLHs, Varaud et al., 2011) and fruit (MADS-boxes, Ripoll et al., 2015). Compound promoter architectures that combine AuxREs with binding sites for other transcription factors would enable specificity and fit well with observed native promoter architectures.

There are many other aspects of auxin signaling that may contribute to specificity in auxin responses, including differential interactions between ARFs and Aux/IAA repressors (Vernoux et al., 2011), differential degradation rate of Aux/IAAs (Havens et al., 2012), and variation in which tissues and at what developmental timepoints ARFs are expressed (Rademacher et al., 2011). As we continue to elucidate the rules of ARF-activated transcription, synthetic tools should make it possible to examine each of these aspects in turn. Future efforts that combine synthetic and native approaches will ultimately be needed to pinpoint the combination of factors that make up the “auxin code”, as well as to make it possible to retrace the evolutionary path that connected novel auxin response modules to diversity in plant form and function.

## Materials and Methods

### Yeast integrating plasmid construction

Oligonucleotides were obtained from Integrated DNA Technologies with standard desalting purification. All cloning was done by Gibson assembly unless otherwise specified, using Phusion high-fidelity DNA polymerase. For yeast constructs, all promoter variant fluorescent reporters were cloned into a URA3-single integrating vector. Promoter variants were ordered as oligo or block gene fragments with Gibson overhangs and cloned the A1 site of a 300 bp IAA19 promoter sequence with a G→A mutation introduced at the second position of each AuxRE site (Pierre-Jerome, et al., 2016). Transcription factors were cloned into a HIS3-targeting single integrating vector under the control of the yeast ADH1 constitutive promoter. Maize ARFs were cloned in pDONR vectors as described in (Galli et al., 2018). After addition of 5’ yeast Kozak sequences (AAA), Gateway cloning (Invitrogen) was employed to integrate ZmARFs into the HIS3-targeting single integrating vector.

### Yeast culturing and transformations

W303-1A yeast cells of mating type locus a (Mata) were cultured in yeast peptone dextrose (YPD), synthetic complete (SC), or synthetic drop out (SDO) media. Media were made according to standard protocols and supplemented with 80 mg/L adenine. Stably integrating constructs were transformed using a standard lithium acetate protocol and plated on selective media plates kept at 30°C. Yeast were glycerol stocked after isostreaking strains on YPD and PCR confirmation of construct integration.

### Flow cytometry assays of ARF activity

A freshly grown colony of each yeast strain was inoculated in 1 mL of SC media and grown at 30°C with shaking at 400 rpm in 2,000 μL Eppendorf Deepwell Plates 96. After 16 hours of growth, cultures were diluted 1:150 into 1 mL fresh SC media. Fluorescence measurements were taken after 4 to 5 hours of additional growth. The data for at least three independently grown replicates were pooled for each strain. Fluorescence measurements were taken with a custom BD Accuri SORP flow cytometer with a CSampler 96-well plate adapter using an excitation wavelength of 514 nm and an emission detection filter at 545/35 nm. A minimum of 10,000 events above a 40,000 FSC-H threshold was measured for each sample. Experiments were done in triplicate for each strain. Data were exported as FCS 3.0 files and processed in R using the flowCore, plyr, and ggplot2 software packages.

## Supplemental Material

Two supplemental figures:

Supplemental Figure S1 Arabidopsis and maize ARFs do not activate on mpIAA19.

Supplemental Figure S2 Arabidopsis ARF7 does not activate on a single AuxRE.

**Supplemental Figure S1 Arabidopsis and maize ARFs do not activate on mpIAA19.** A) Activity of AtARF5 and AtARF19 on the mpIAA19 promoter, with all the AuxREs mutated. B) Activity of ZmARF4, ZmARF27, and ZmARF29 on the mpIAA19 promoter, with all the AuxREs mutated.

**Supplemental Figure S2 Arabidopsis ARF7 does not activate on a single AuxRE**. Despite a conserved asparagine shared with AtARF19 within the DD domain, AtARF7 does not activate on a single AuxRE of the sequence 5’-TGTCGG-3’, but activates on two AuxREs of this sequence facing each other.

## Acknowledgements

Thank you to members of the Nemhauser and Imaizumi labs for helpful discussion and guidance on experimental design and execution. Thank you especially to Manraj Sahota, Mollye Zahler, and Arjun Khakhar for initial experimental work and many discussions. This work was supported by the National Science Foundation (MCB-1411949), and National Institute of Health (R01-GM107084) and the Howard Hughes Medical Institute Faculty Scholar Award. AL was supported by an NSF Graduate Research Fellowship DGE-1256082. MMTT was supported by an NSF Postdoctoral Fellowship in Biology IOS-1609014.

## Author Contributions

Experimental design was conceived by AL, MMTT, and JLN. Research was performed by AL, MMTT, and EAO. The manuscript was prepared by AL, MMTT, and JLN.

## One-sentence summary

The plant growth hormone auxin regulates development via a family of transcription factors that share promoter sequence preferences, despite activating different genetic networks.

